# The Olfr151 Odorant Receptor Gene is Resistant to Activation in Embryonic Stem Cells

**DOI:** 10.64898/2026.02.19.706738

**Authors:** Irena Parvanova, Eugene Lempert, Paul Feinstein

## Abstract

A single allele of any of the ∼1100 functional mouse odorant receptor (OR) genes is expressed by mature olfactory sensory neurons (mOSNs) through a poorly understood mechanism of singular gene choice. We were interested in developing an expression system to study OR selection in a mouse embryonic stem cell (mESC) carrying an Olfr151-IRES-CRE knock-in and a CRE-recombination-dependent ROSA26-MTMG reporter. Recombination of the ROSA reporter could not be observed after exposing this mESC to thousands of chemical compounds, including 175 compounds known to interfere with epigenetic regulatory processes nor by transfection with high probability Olfr151 promoter minigenes coexpressing CRE. The only two known regulatory elements that control OR promoters are not olfactory specific. Still, our inability to elicit even leaky CRE recombination in mESCs suggests that a specific transcriptional machinery within the OSN lineage is needed to drive OR promoter activation.

**Author Summary:** The Olfr151 promoter is inactive in mESCs.

## Introduction

Ten million olfactory sensory neurons (OSNs) are arrayed in the main olfactory epithelium (MOE), located inside the nasal cavity. The mucosal layer facing the luminal side of the MOE is densely populated with ciliary extensions originating from OSNs. These cilia contain the odorant receptors (ORs) capable of binding odorant molecules (ligands)(*1–4*). ∼1100 functional OR genes belong to the well-characterized G-protein-coupled receptors (GPCRs) superfamily (*5, 6*). While many non-chemosensory GPCRs are readily expressed in heterologous cells, ORs have had difficulty being analyzed *in vitro* (*5, 7*). Additionally, the apparent singular expression of ORs in the olfactory epithelium, where one allele is expressed per OSN, means that each allele has only ∼4500 OSNs associated with its expression (*8, 9*). Understanding the mechanism of singular OR expression will allow for the isolation and characterization of the singular gene choice transcriptional complex common to all OSNs (*10*).

According to the prevailing model of singular OR expression, all OR genes within the postmitotic immature neurons are rendered in an inactive state (transcriptionally silenced), which is followed by one or several OR genes being expressed. OR genes are believed to be actively silenced through the addition of repressive epigenetic chromatin modifications, such as H3K9me3 and H4K20me3 (*11*). This model suggests several OR alleles initially have their repressive histone modifications replaced with H3K4me3 modification, a hallmark of transcription initiation. Only one allele “wins” that makes the most mRNA through its association with one of the solitary super enhancers, aka a Greek Island Hub. Much of the initial conceptual framework stems from work by Magklara et al. (*11*), which observed increases in H3K9me3 and H4K20me3 repressive epigenetic modifications on the promoters of the OR genes within mature OSNs compared to non-olfactory, Liver cells. These data suggested that chromatin modifications that silence OR genes are absent in liver cells. Thus, the increase in the repressed status was interpreted as the manifestation of an OR singular gene choice mechanism that is initiated by silencing OR genes in the OSNs (*11*).

A previous analysis of embryonic stem cells (ESCs) showed that over 75% of the genes are marked with H3K4me3 transcription initiation chromatic epigenetic mark, even if the genes are not expressed (*12*). But OR genes were among the 25% not marked with H3K4me3 transcription initiation epigenetic mark (*12*); They may also be marked with the repressive epigenetic chromatin marks (*12*). A broader interpretation of these findings is that OR genes are indeed marked by H3K9me3 and H4K20me3 in most cell types and this repressive state increases in olfactory neurons but is not necessarily linked to the mechanism of selecting one OR allele for expression (*12–17*).

Since the identification of OR genes, it has only been possible to study their gene regulation within an animal model, where OR mRNAs are robustly produced, and the translated OR protein traffics to the plasma membrane in both olfactory cilia and along axonal processes (*8*). Many years ago, we showed that double heterozygote mice with the bicistronic Olfr151-IRES-CRE (recombinase) knockin and a CRE reporter would faithfully mimic the expression of the bicistronic Olfr151-IRES-GFP in OSNs (*18*). Unfortunately, no cell line exists that stably expresses ORs in an OSN-type precursor through the mechanism of singular gene choice. Therefore, we attempted to develop an *in vitro* assay by forcing mouse embryonic stem cells to activate OR gene expression.

As a first step to create a cell line that could express OR genes, we generated a mESC line by crossing the Olfr151-IRES-CRE mouse line with a ROSA26 reporter mouse line (*18, 19*). In this ROSA26 mouse line, a membrane-anchored tdTomato fluorescent (MT) coding sequence, located between two loxP sequences, is followed by a membrane-anchored GFP (MG) coding sequence (*20*). Subsequently, a mouse embryonic stem cell line was generated from Olfr151-IRES-CRE; ROSA26-MTMG blastocysts. If the Olfr151 gene is expressed in this new mESC line, then the ROSA26 gene locus should switch to express MG (*21*) [CRE would excise the membrane tdTomato (MT) sequence, located between two loxP sequences, allowing for green fluorescence to be observed]. Because Olfr151 is not expressed in mESCs, no MG expression was observed. We will refer to this cell line as Olfr151iCRE; ROSA26-MTMG (i=IRES) mESCs in the rest of the manuscript.

We attempted to force the Olfr151 OR gene expression by setting up a high-throughput small compound chemical library screen assay. We administered approximately 5000 compounds to Olfr151iCRE; ROSA26-MTMG mESCs, including 175 known to interfere with epigenetic processes. Per our observation, none of the chemical compounds were able to cause a red to green fluorescence shift in the mESCs. Perhaps the absence of CRE expression was due to the poorly activated Olfr151 promoter. To increase the probability of Olfr151 expression, a strong singular gene choice enhancer (5×21) is be added, which radically increases its expression as we observe in 5×21-Olfr151iCRE; ROSA26-MTMG mice. Using our Olfr151iCRE; ROSA26-MTMG mESCs, we transiently expressed a 5×21-Olfr151iCRE minigene (*22*) and exposed these cells to the 175 epigenetic compounds. Despite our efforts, we failed to observe expression of our reporter in mESCs.

## Materials and Methods

### High-throughput screenings assay, using small chemical compound libraries

Blastocyst-derived mESCs were isolated from Olfr151iCRE;ROSA26-MTMG mouse line (*21*). The cells were plated onto 384-well plate (Corning®, cat. #3683), with the seed density of 0.3 x10^4^ cells/well. In 24h, the cells were treated with chemical compounds from LOPAC (1280 compounds, Sigma-Aldrich-Millipore, Lopac®1280), (Prestwick chemical library (1280 compounds Prestwick Chemical, provided in DMSO), and MicroSource library (2300 compounds, MicroSource Discovering System, Inc., provided in DMSO), at 1µM and 10µM concentration.

The Olfr151iCRE;ROSA26-MTMG cells with the chemicals were grown at 37°C at 5% CO_2_ overnight. The next day, the cells were fixed in 1% paraformaldehyde solution EM Grade (PFA, Electron Microscopy Sciences) and washed three times in 1X Phosphate Buffered Saline (PBS, Corning). The cells remained in PBS until imaging.

Data were analyzed at the Cell Screening Core of Weill Cornell Medical College, using ImageXpress MICRO imaging system from Molecular Devices equipped with a Photometrics CoolSnapHQ camera from Roper Scientific. tdTomato images were acquired using 543/22 nm excitation and 593/40 nm emission filters with a 569 nm-dichroic long-pass filter. GFP images were acquired using 472/30 nm excitation and 520/35 nm emission filters with a 502nm-dichroic long-pass filter. Plates were transported from plate hotels using a CRS CataLyst Express robot (ThermoFisher Scientific). Overall, images were acquired at four separate sites per well. In two instances, nine sites were acquired per well. The imaged sites were selected at the center of the well with 200µm spacing between sites. 696 x 520 pixel images (897 x 670µm) were acquired at 12 intensity bits per pixel. Each pixel is 1.29 x 1.29µm in the object. The values were provided by Weill Cornell Medical College Screening facility.

### ChIP-qPCR assay to investigate epigenetic marks on ORs in mESCs

Olfr151iCRE; ROSA26-MTMG mESCs were grown on 10cm (100×20 mm Tissue Culture Dish, Corning), at confluency of 30-45 x 10^6^ cells. The cells were fixed with PFA (PFA, Electron Microscopy Sciences), added to the cell growth media at a final concentration of 1% PFA at 37°C for 10min. Incubation was followed by quenching the reaction with glycine, added to the cell growth media at a final concentration of 0.125M for 5min at RT (shaker). Cells were washed 4 times with cold 1X Phosphate Buffered Saline (PBS, Corning) and scraped from the plate surface. For this purpose, we used 1mL ice-cold 1X PBS (4 x 1ml). We collected them into 15 mL conical tube (15 mL Centrifuge Tube, CELLTREAT). Cells were spun down at 2000 rpm for 15min at 4°C. The cells were frozen at −80°C before the next step.

The cells were resuspended in RIPA buffer (Sigma) with PIC (cOmpleteTM, Mini, EDTA-free, Roche) and PMSF (32%, Electron Microscopy Sciences) (2.5 mL RIPA buffer/ 50 x 10^6^cells). The cells were incubated on ice in 1X RIPA buffer (Sigma) for 30min. To lyse the cells, we vigorously vortexed them every 5min (the cell lysis can be checked under a microscope). 1 mL aliquots of the samples were transferred to microcentrifuge tubes. To shear the DNA, the cell lysate was sonicated in a cup-horn digital sonicator (on ice). 1min sonication at high amplitude was followed by 1min rest on ice (30 times). Cells were spun down at 13,000 rpm for 20min at 4°C. The supernatant was transferred into new tubes. If there was more than one aliquot for the same 10cm plates (100 x 20 mm Tissue Culture Dish, treatment, the samples were pooled. Aliquots were made and stored at −80°C (*23*).

### BCA assay was done for protein concentration (Pierce® BCA Protein Assay, Pierce)

40µg protein (10%) was used as input, 800µg protein was used for immunoprecipitation (IP) (400µg aliquot is used for IP, 400µg aliquot is used for a negative control with a non-specific antibody, IgG), and 100µl (or less) for DNA shearing check.

400µg of protein was pretreated with Protein A/G PLUS-Agarose beads (Santa Cruz Biotechnology) for 30min at 4°C. Once the beads were removed, we added an antibody of interest to the protein sample as follows:

- H3K9me3 (Anti-Histone H3 (tri methyl K9) antibody – ChIP Grade ab8898, Abcam),
- H4K20me3 (Anti-Histone H4 (tri methyl K20) antibody – ChIP Grade ab9053, Abcam),
- H3K4me3 (Anti-Histone H3 (tri methyl K4) antibody (mAbcam1012) - ChIP Grade ab1012, Abcam),
- IgG (negative control) (Normal rabbit and mouse IgG, Santa Cruz Biotechnology).

The optimal concentration of each antibody can vary; therefore, an Antibody (Ab) titration curve was performed. Based on our result (data not shown), we used 6µg of the treatment Ab. The samples with added Abs were incubated overnight at 4°C. Next day, we used Protein A/G PLUS-Agarose beads (Santa Cruz Biotechnology), pre-blocked with 0.3mg/mL of salmon sperm DNA for 30min at 4°C to pull down the Ab-bound proteins. blocked the beads. We washed the beads 3 times with cold 1X PBS by spinning them down for 2min at 3000rpm at 4°C. After the wash, approximately 25% of bead slurry remained in a50µl beads aliquot. 50µl bead slurry was added to the immunoprecipitation (IP) sample (a cell lysate + antibody) and incubated for 2h at 4°C.

After 2h, the samples were spun down for 2min at 3000rpm at 4°C. The supernatant was removed. The samples were washed with 1ml of the following solutions (washing solutions components are included) and spun for 2min at 3000rpm at 4°C between washes: Wash 1, Wash 2, Wash 3, TE buffer twice (pH=8).

After the washes, 100µl of 1X Elution buffer (freshly made)/5µl of Proteinase K (20mg/mL, ThermoFisher Scientific) were added. 10µl of 10X Elution buffer/5µl of Proteinase K (20mg/mL) were added to the samples. For input, we adjusted the volume of the 40µg aliquots of untreated sample to 80µl using 1X RIPA buffer. 11µl of 10X Elution buffer/5µl of Proteinase K (20mg/ml) were added to the DNA shearing sample.

### ChIP Wash Solutions

For 500µl final volume, adjust volumes with H_2_O and filter:

Wash 1: 0.1% SDS (5mL of 10% SDS)

1% Triton-X (5mL of Triton-X 1000)

20mM Tris pH8 (10mL of Triton-X 100)

150mM NaCl (15mL of 5M stock solution)

Wash 2: 0.1% SDS (5mL of 10% SDS)

1% Triton-X (5mL of Triton-X 100)

20mM Tris-Cl pH8 (10mL of 1M stock solution)

500mM NaCl (50mL 5M stock)

Wash 3: 0.25M LiCl (5.3g LiCl mol. Weight 43.39g/mol)

1% IGEPAL/NP-40 (5mL)

1% Deoxycholate – 5g NaDeoxycholate

1mM EDTA (1mL of 0.5M stock solution)

10mM Tris pH8 (5mL of 1M stock solution)

10X Elution Buffer preparation

10X Elution Buffer was kept at 65°C to avoid viscosity. For 500µl buffer, we added 10% SDS and 0.042g sodium bicarbonate (NaHCO3, Sigma). We used H_2_O for 1X dilution. After adding the elution buffer, the sample tubes, the input, and the sheared DNA sample were incubated at 65°C overnight. On the following day, DNA sample was purified, using a QIAGEN PCR Purification kit. qPCR was used to analyze the results.

The qPCR primers were initially applied in a regular PCR (dNTPs, Invitrogen; Titanium Taq Polymerase, Clontech Laboratories), using input DNA as a template. We additionally prepare and run an agarose gel electrophoresis to assess the presence of a product. The qPCR primers were tested on various dilutions of genomic DNA template from Olfr151iCRE; ROSA26-MTMG cells to determine their efficiency(*24*).

### Primer sequences

GAPDH (*25*)-*gapdh* (4333764) from Applied Biosystems Assays on Demand primers 5’-AGGTGAAAATCGCGGAGTG-3’ & 5’-AGCATCCCTAGACCCGTACA-3’

β-globin (*26*)

5’-GGACAGGTCTTCAGCCTCTTGA-3’ & 5’-CAGATGCTTGTGATAGCTGCCT-3’

Olfr151 promoter (designed by us):

5’-TGATAAGGATGGGTGGGTAGT-3’ & 5’-TAGCATCAACAGAGCCGTTAC-3’

Primer efficiencies found in Supplemental Table 1.

### Wholemount olfactory bulb imaging

We imaged wholemount olfactory bulbs with LMS510 laser scanning confocal microscope using a 5X objective. GFP was excited at 488nm and collected at BP500-545nm, while tdTomato was excited at 561nm and collected at LP575nm.

## Results

### Olfr151iCRE expression in the olfactory system

Approximately 1100 odorant receptor (OR) genes are expressed in OSNs in a monoallelic and monogenic fashion. Therefore, approximately 2200 OR alleles are singularly expressed in OSNs. As such, the addition of an odorant receptor transgene to the mouse genome leads to an extra allele for this process of singular gene choice. Knockin mice for Olfr151 are expressed in a few thousand OSNs.

We have previously reported the insertion of IRES-CRE (iCRE) into the odorant receptor Olfr151 locus (Olfr151iCRE) crossed to the Z/EG mouse reporter [actin promoter driving a floxed (flanked by same orientation loxP sites) LacZ followed by GFP]. Olfr151iCRE; Z/EG labeled Olfr151 axons with GFP and coalesced with Olfr151-IRES-TauRFP_2_ (*18*). A schematic of the ROSA26 locus is shown in Fig. 1A, which also carries an actin promoter to drive a floxed membrane-bound tdTomato fluorophore (MT) followed by membrane-bound GFP (MG) (ROSA26-MTMG). The activation of the Olfr151 gene, leads to the expression of Olfr151 OR and CRE proteins (*18*). As expected, a single green focal point of coalesced axons (glomerulus) from OSNs expressing Olfr151 is observable in each hemisphere of the dorsal olfactory bulb (Fig. 1B).

**Fig. 1.**
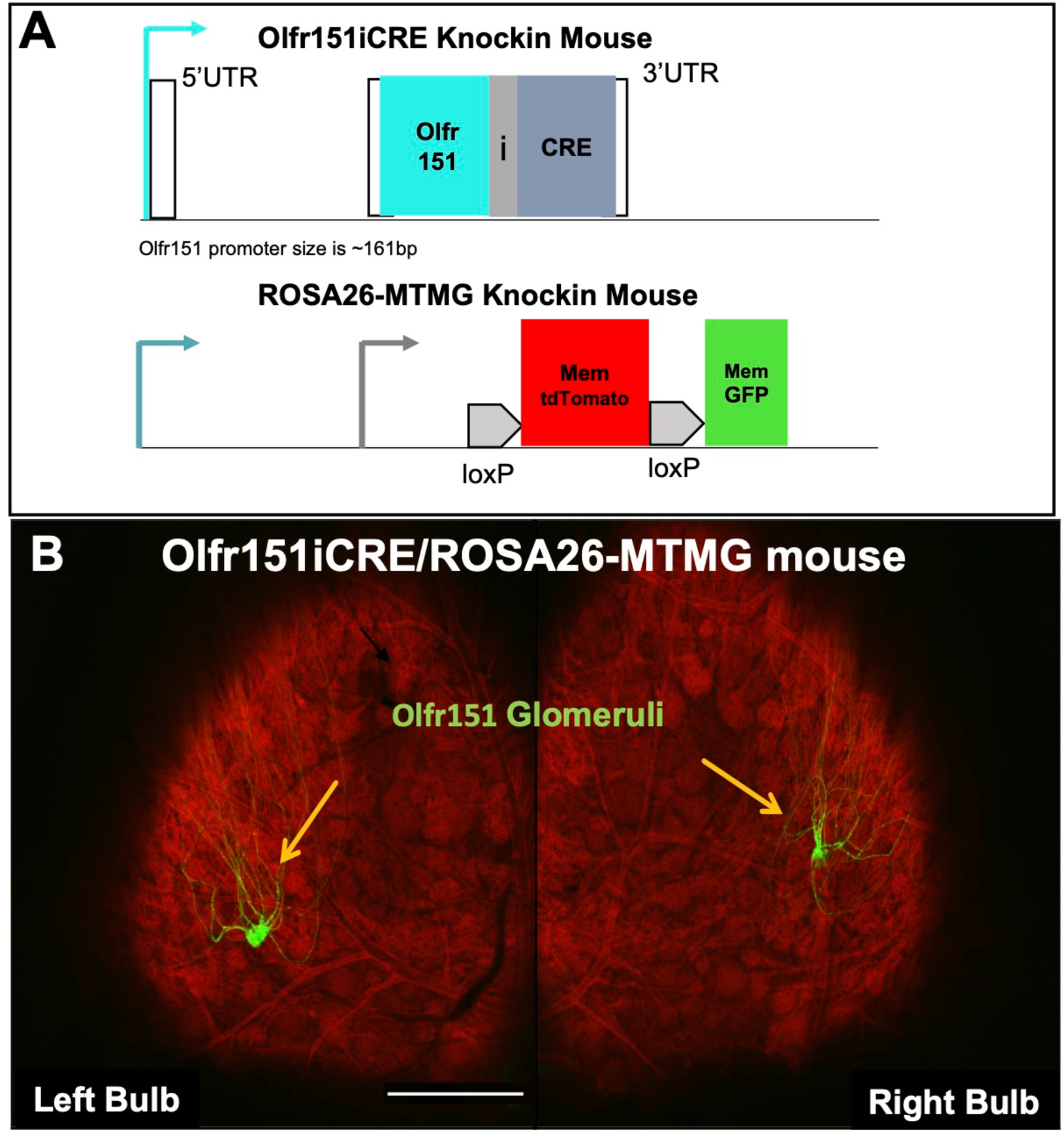
Olfr151iCRE; ROSA26-MTMG double heterozygous mouse (schematic and confocal microscope image). **(A)** IRES-CRE is inserted in the Olfr151 OR locus on chromosome 9 (Olfr151iCRE). The loxP-Membrane (Mem)-tdTomato-loxP-Mem-GFP (MTMG), CRE-Reporter construct is gene targeted into the ROSA26 locus on chromosome 6 (ROSA26-MTMG). **(B)** Confocal wholemount fluorescence microscopy of mouse olfactory bulbs in Olfr151-IRES-CRE mice; ROSA26-MTMG. All cells in the bulb reveal red fluorescence except axons derived from Olfr151iCRE expression, which are green and coalesce into the depicted green glomeruli (yellow arrows). Scale bar =200µm.

### Can the Olfr151 promoter be activated in embryonic stem cells (ESCs)?

Decades of effort into immortalizing progenitors of OSNs have led to immortalized olfactory cell lines that do not reveal stable OR mRNA expression (ODORA, OP6/OP27, 2.2.1/ 1.1.17)(*27–29*). We undertook a new approach to activate OR gene expression utilizing embryonic stem cells as a self-renewing progenitor population. In this new effort, an embryonic stem cell line was derived from double heterozygous Olfr151iCRE; ROSA26-MTMG blastocysts (*21*). Characterization of this mESC line revealed three critical features. First, Oflr151iCRE knockin is silent as ESC colonies readily expressed membrane-tdTomato at the plasma membrane (PM), but were devoid of any membrane-GFP expression due to the absence of any leaky or stochastically expressed CRE protein from the Olfr151 locus (Fig. 2)(*18*). Second, OCT3/4 protein, a hallmark of ESCs is robustly expressed (Fig. 2D). Finally, the derived mESC line formed colonies and propagated like typical mESCs, with doubling times of less than 1 day. In addition, karyotyping results for 54 of these ESCs show 69% normal counts and 31% with Trisomy 1. To confirm the functionality of the ROSA26-MTMG reporter, a plasmid with CMV promoter-driven CRE (pOG231, a gift from Geoff Wahl, Addgene plasmid #17736) was transfected into Olfr151-IRES-CRE;ROSA26-MTMG mESCs (*30*). When an exogenous source of CRE is added to these ESCS, mosaic expression of membrane GFP fluorescence is observed (Fig. 2E-G). GFP expression at the plasma membrane is clearly shown under high magnification (Fig. 2H). Thus, a new mESC line has been derived carrying a functional CRE reporter with a silent Olfr151iCRE OR locus.

**Fig. 2.**
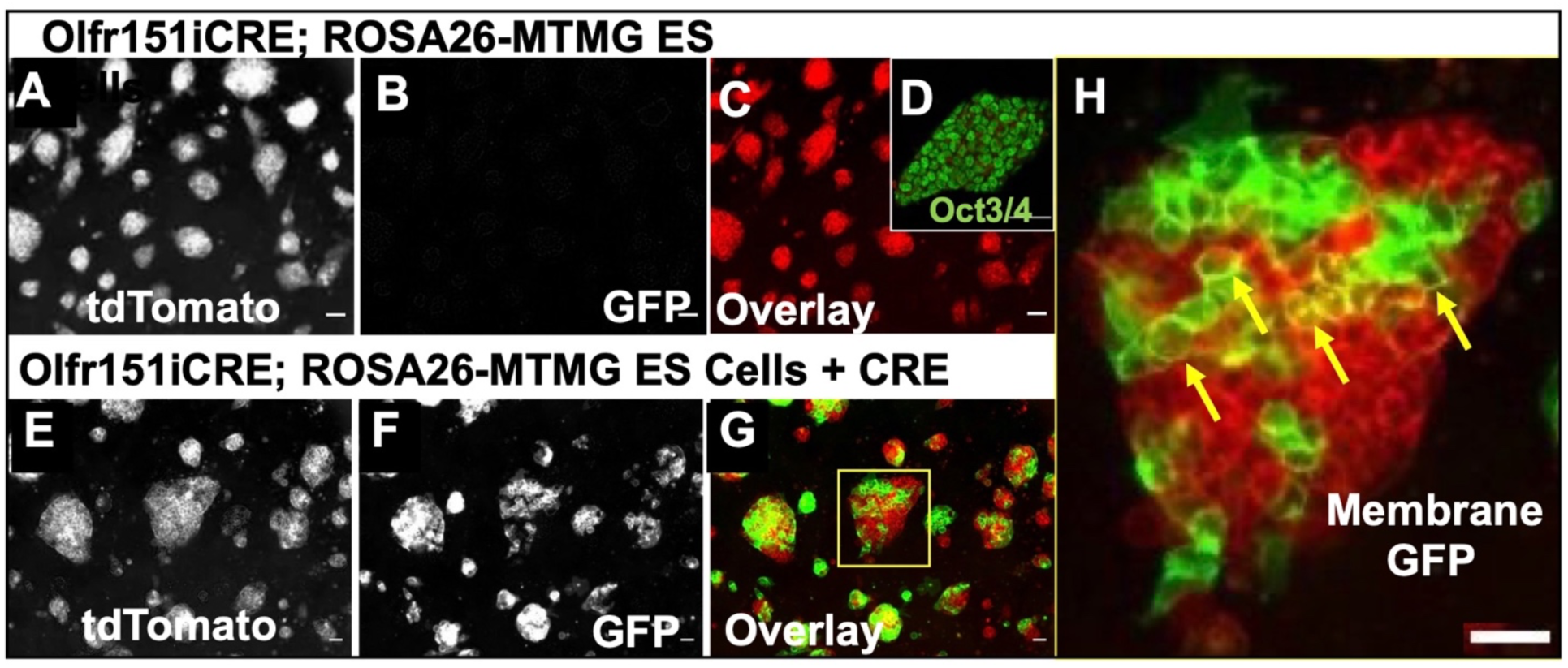
Olfr151iCRE; ROSA26-MTMG derived embryonic stem cell line confocal microscope images. **(A-H)** Olfr151iCRE; ROSA26-MTMG mESCs were derived from blastocysts of the Olfr151iCRE; ROSA26-MTMG mouse line. **(A)** All blastocyst-derived mESCs emit red fluorescence due to the presence of the tdTomato expression at the cell membrane. **(B)** No membrane GFP fluorescence is observed, suggesting the absence of leaky CRE expression from the Olfr151 locus. **(C)** Overlay image of tdTomato and GFP channels expression of transfected mESCs. **(D)** Oct3/4 immunofluorescent (IF) staining of Olfr151iCRE; ROSA26-MTMG mESCs labels all cell nuclei. Scale Bar =20µm (E-G) Images of Olfr151iCRE; ROSA26-MTMG mESCs transiently transfected with pOG231(nls-CRE). **(E)** All blastocyst-derived cells mESCs emit red red fluorescence, which reflects the presence of the tdTomato expression at the cell membrane. **(F)** GFP fluorescence is observed reflecting CRE expression at the mESC membrane. **(G)** Overlay image of tdTomato and GFP channels expression of transfected mESCs. **(H)** High magnification of a cell colony shows membrane GFP localization (yellow arrows). Scale Bars =100µm.

### High-throughput screening platform utilized to activate Olfr151 gene in mESCs

Small chemical molecules have been used to identify self-renewal properties in mESCs (*31*). Libraries of these chemical compounds allow for the establishment of high-throughput assays in 384-well plates, a standardized, cost-effective, and stable platform. Here, approximately 4860 screening compounds were exploited, at two different concentrations (1µM and 10µM) from LOPAC chemical library by Sigma Aldrich: ∼1280 compounds, Prestwick library by Prestwick Chemical, which contain ∼1280 compounds, and MicroSource by MicroSource Discovering System, Inc.: ∼2300 compounds (Fig. S1). The selected concentrations were picked based on standard practice in small compound libraries treatment analysis (*31*). If CRE expression were to be activated from the Olfr151 locus, then membrane-GFP fluorescence would be observed.

### Setting Up High-throughput Screen

Can CRE-mediated events in the Olfr151iCRE; ROSA26-MTMG mESCs in 384-well plate be visually identified? Our mESC line, tdTomoto fluorescent mESC colonies are readily observed, lacking green membrane fluorescence. The data were quantified as log10 of tdTomato fluorescent area/Area A in µm^2^ and GFP fluorescent area/Area B, µm^2^, based on the mESC image readouts by ImageXpress MICRO imaging system (Fig. 3A, B). Up to nine images could be taken per well and quantified corresponding to 75% of the well surface area. Four images taken per well constitute 34% of the well area. Our analysis of the numerical values of red and green fluorescence (log10), reveal the average tdTomato fluorescent baseline value for untreated Olfr151iCRE; ROSA26-MTMG mESCs is log10_avg_ = 5.77 ± 0.09 µm^2^ (x-axis), while the GFP fluorescent baseline is log10_avg_ = 2.77 ± 0.37 µm^2^ (y-axis) (Fig. 3C-E). Adding a standard deviation value of 0.37 to the highest baseline GFP value of 3.14 (2.77 ± 0.37) in untreated mESCs gives us maximum baseline of 3.51 µm^2^. By contrast, when a CRE expressing plasmid was transfected and mESCs plated on 384-well plates, both tdTomato and GFP expression were apparent by visual inspection (Fig. 3D). Quantification of GFP expression in mESCs with transiently transfected CRE plasmid now shows GFP fluorescence is log10_avg_ = 4.84 ± 0.16 µm^2^ (y-axis), which is well above the established background GFP baseline (Fig. 3E) (*31*).

**Fig. 3.**
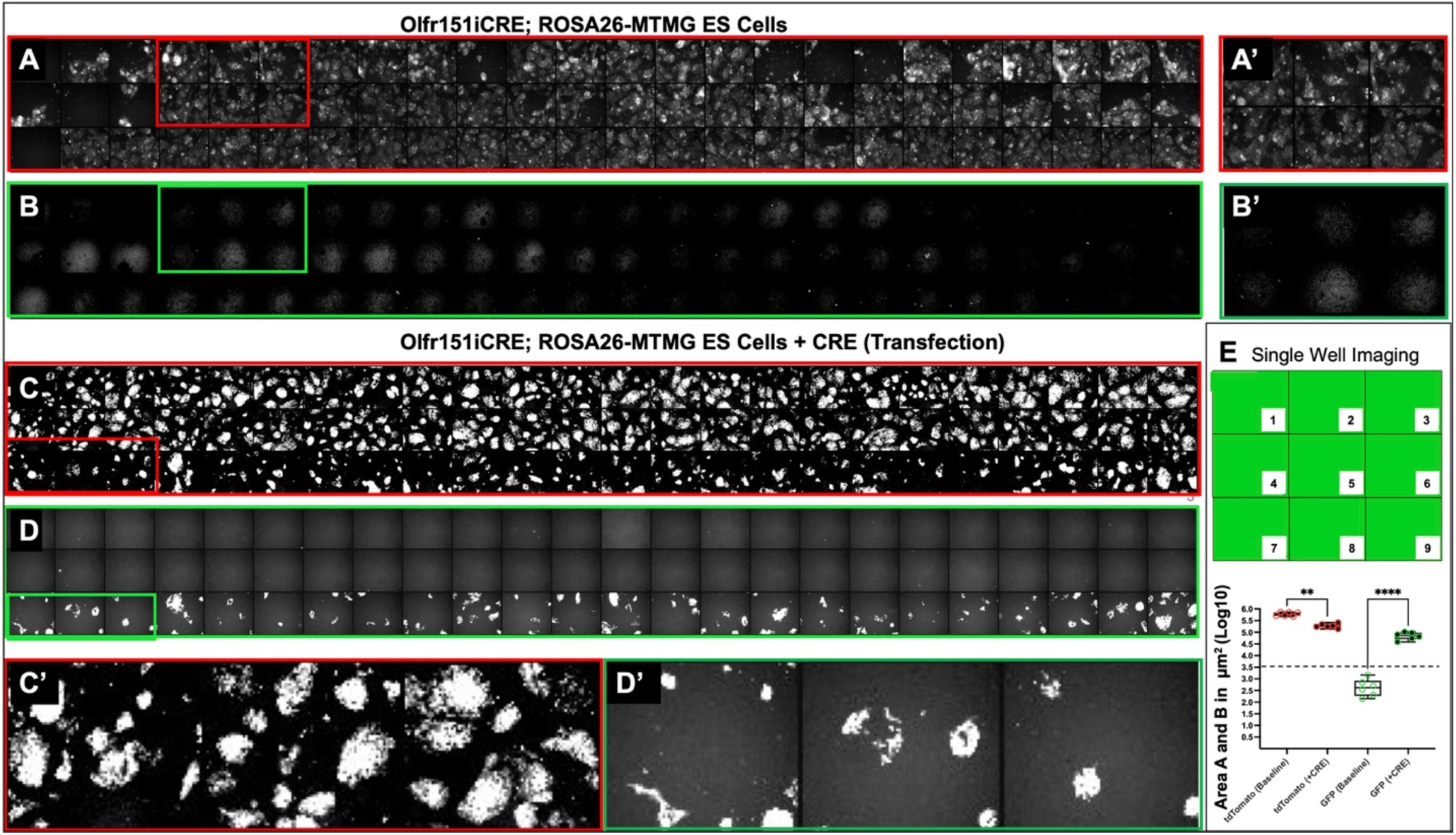
Confocal images of Olfr151iCRE; ROSA26-MTMG mESCs transfected with CRE plasmid. (A) Robust tdTomato fluorescence expression is observed in untreated mESCs from 72 wells, which include the first three rows of a 384-well plate). (B) No GFP fluorescence is observed in these 72 wells. (A’, B’) Zoomed-in images for 6 /72 well noted above. (C) We observed robust red fluorescence in the first two rows of untreated cells (48 wells) and in the one row (24 wells) of mESCs transfected with CRE plasmid (pOG231). (D) We only observed robust green fluorescence in the third row of the CRE transfected cells. In 24 hours, the cells were fixed and imaged. (C’, D’) Zoomed-in images for 3 wells. (E) Up to nine images were taken per well, representing 75% of the well surface area. If four images are taken per well, 34% of the well area is captured. (F) The fluorescence values are expressed as log10 Area (fluorescence area, µm^2^). tdTomato and GFP baseline values (background fluorescence) depicted. We determined the 3.5µm^2^ value (y-axis) by adding the maximum value of GFP baseline from the log10 Area mean of 3.16µm^2^ in the untreated mESCs and a standard deviation of 0.37. Any value over 3.5µm^2^ could potentially signify CRE expression in the reporter (****p<0.0001, ANOVA; error bars indicate mean +/-SD). By contrast GFP (CRE+) had the lowest value of 4.59 ± 0.16µm^2^, which is well above the baseline max value of 3.16 ± 0.37µm^2^.

### Library Screens

High-throughput screens were performed for all three chemical library compounds. The first screen utilized the smallest chemical library (LOPAC) in a single concentration of the chemical. mESCs in each well of a 384-sample plate were treated with a different chemical (Fig. 4A-C). Since the LOPAC chemical library contains ∼1280 compounds, we treated four 384-well with plated mESCs with a single chemical per well at 1µM. The final row (O) on each plate was left empty (no mESCs were plated) and not treated with a chemical to quantify background autofluorescence in the red and green-fluorescent channels.

**Fig. 4.**
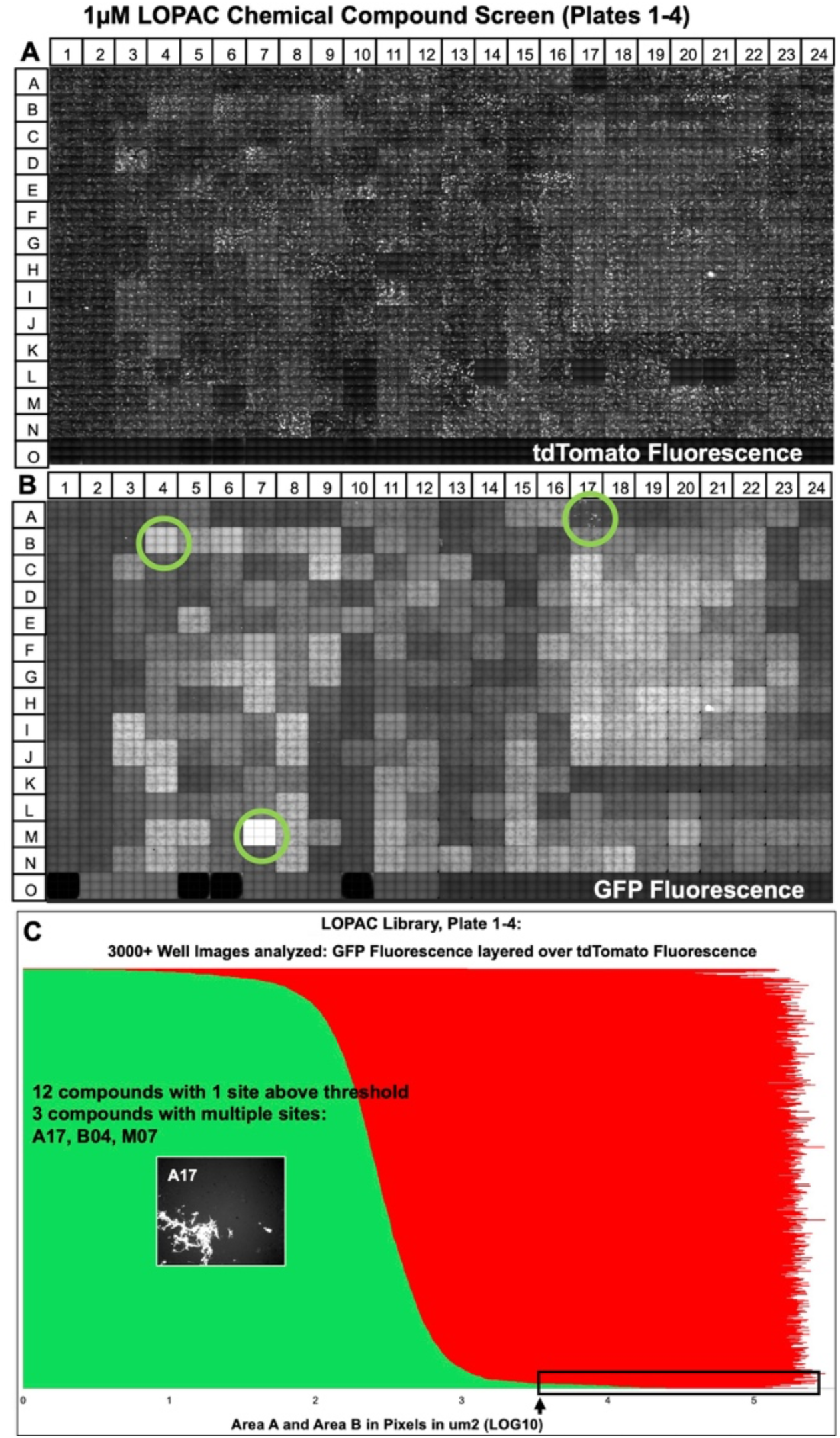
An example of fluorescent readings after LOPAC chemical compounds treatment of mESC reporter cell (384-well plate setup). Reporter cells were plated at low density. In 24h, the cells were treated with LOPAC chemical compounds at 1µM concentration. (A) Grayscale of 9 images per well were acquired at tdTomato fluorescent frequency (543/22nm excitation; 593/40nm emission filters). **(B)** Grayscale for 9 images per well were acquired for GFP (472/30 nm excitation; 520/35nm emission filters). (**C)** tdTomato and GFP fluorescence expression (Log10 fluorescence area, µm^2^) in reporter cells after LOPAC compound treatment. 34 of the captured images exhibit GFP fluorescence above 3.5µm^2^ (threshold fluorescence; see arrow and rectangular box). GFP fluorescence were captured in a 12-well plate were treated with a single compound per well above threshold. GFP fluorescence above the threshold in three wells treated with a single compound per week was identified: A17, B4, M7 (green circles on wells). Autofluorescence is observed in wells B4 and M7. Autofluorescence crystals that do not follow membrane GFP morphology are found in well A17 (see inset). See Supplemental Fig. S2 for all LOPAC screenings yielding 6881 images (for the remainder of the plates, we took 4 images/well). In 123 images, the GPF fluorescence was above log10 Area B = 3.5µm^2^.

The imaging data were acquired 48h after the chemical treatments. Initially, 9 composite images per well were collected (∼75% of the surface area) for the mESC treated with LOPAC small compounds. tdTomato fluorescence show punctate patterns of mESC colonies (Fig. 4A) as previously observed in Fig. 3A, C. GFP fluorescence was diffuse (Fig. 4A). Only wellA17 exhibited punctate green fluorescence (Fig. 4B). No autofluorescence in the red (tdTomato) emittance channel was observed, as shown in row O (Fig. 4A). Conversely, GFP autofluorescence was readily observed in the green (GFP) emittance channel in row O of the 384-well plate where no mESCs were plated (Fig. 4B). Quantification from over 3000 red and green fluorescence values were overlayed and ranked from least green fluorescence to most (Fig. 4C). It should be noted that red fluorescence values predominantly from tdTomato with log10_avg_= 5.00±0.55 did not vary significantly from mESCs red fluorescent values in the control images in Fig. 3E (log10_avg area_ values = 5.77 µm^2^). From this initial screen of four 384-well plates depicted in Fig. 4C, we identified 34 images among 12 wells (chemicals) with mESCs that had GFP fluorescent value of log10_area_ > 3.50. 7 images were taken in wells numbered A17, B04, and M07. But, none of the wells showed membrane-bound GFP expression. For instance, in Fig. 4C, the punctate GFP expression in the chemically treated mESC in well A17, resulted from the crystals from the compound itself. In a subsequent analysis, we visually examined 6681 images of mESCs treated with LOPAC small compounds library at concentrations of 1µM and 10µM with 4 images captured per well. In total, 123 well images were identified with log10 values > 3.50 (Fig. S2). mESC colony morphologies were inspected in each image from the 123 images. If a brief event of OR expression took place, then this might have resulted in just a few cells in the colony with membrane GFP expression. In addition, several chemicals were examined that were just below the log10 _area_ values of 3.50. All three library screenings resulted in a list of 455 chemicals that were potential hits in activating Olfr151 promoter in mESCs. Autofluorescence was observed for 400 of the potential chemicals, leaving 55 compounds log10 _area_ values > 3.50 (Fig. 5A) to be rescreened at concentrations of 1µM and 10µM and inspected for localization of membrane-GFP. None of the images revealed membrane expression as shown in Fig. 2H. A representative image is shown for four chemicals, which show particulates generating autofluorescence (Fig. 5B-E). One of these four chemicals, piperacetazine shows a few regions with isolated fluorescence, but on closer inspection, the fluorescence is not localized at the membrane. In sum, none of the 4860 chemical compounds, from all three libraries screened at two concentrations, showed any promising effects on the activation of the Olfr151iCRE reporter in the mESCs.

**Fig. 5.**
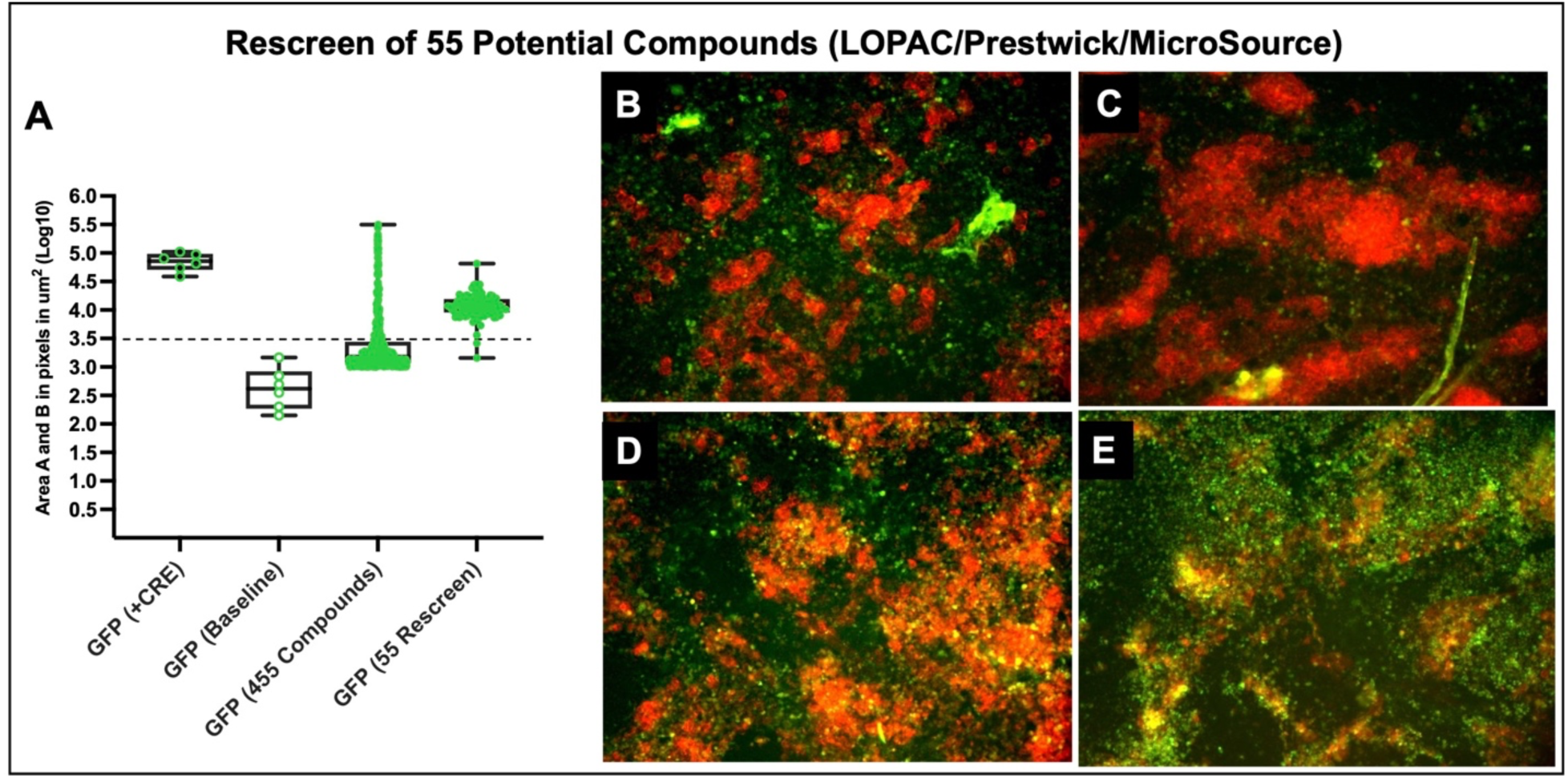
We used three separate small chemical compounds libraries in two concentrations (including LOPAC) to treat our reporter mESCs. **(A)** LOPAC, Prestwick, and MicroSource libraries compounds were used to treat mESCs (at 1uM and 10uM concentration), yielding 455 candidate compounds (log10 Area B Mean value = 3.37µm^2^). A systematic review of the images generated a list of 55 compounds (log10 Area B Mean value = 3.73µm^2^) to be rescreened that were significantly above the log10 Area 3.5µm^2^ threshold (see Supplemental Table 2). **(B-E)** Four examples of compounds that were used for the mESC treatment (rescreened compounds) showed autofluorescence or/and were lackingcellular morphologies consistent with membrane GFP localization. **(B)** Docetaxel (a chemotherapeutic drug) screening results. **(C)** Piperacetazine (an antipsychotic drug) screening results. **(D)** Metrizamide (Non-ionic contrast medium compound) screening results. **(E)** (-)-Eseroline fumarate (a potent analgesic) screening resuluts. Scale bar = 100µm. **(B-D)** The compound treatment in these panels was done in lower compound concentration (1µM) and (E)Compound concentration treatment at 10µM.

### H3K4me3 and H3K9me3 marks for the Olfr151 OR locus in derived mESCs

We were intrigued by the Guenther et al. studies on human embryonic stem cells (hESCs) where the authors showed that the H3K4me3 transcriptional initiation marks are absent in hESCs for OR loci despite being found in 75% of all other genes in the model system (*12*). To confirm and extend this analysis, we performed a chromatin immunoprecipitation-quantitative polymerase chain reaction assay (ChIP-qPCR) to determine the H3K4me3 epigenetic chromatin landscape on Olfr151 OR promoter in mESCs. A comparison to the GAPDH gene expression was made (a gene known to be expressed in mESCs). Conversely, beta-globin gene (hbb) was used as a reference for a gene that is expressed later in development in a cell-type-specific manner (*32*). The presence of H3K9me3 negative epigenetic marks was analyzed for all three genes as well. The ChIP-qPCR analysis were performed in triplicates during three separate experiments. The H3K4me3 immunoprecipitation (IP) readings, normalized to control IgG, revealed the amplification of GAPDH and the lack of amplification for Olfr151 and hbb sequences (Fig. 6A). By contrast, the H3K9me3 off marks IPs for all three genes showed some degree of amplification. (Fig. 6B). Using one-way ANOVA statistical analysis, we confirmed a greater than 30-fold enrichment (*p<0.05) for the presence of H3K4me3 mark at the GAPDH promoter in comparison to Olfr151 and hbb genes. Not surprisingly, the ratio of H3K4me3/H3K9me3 for GAPDH compared to Olfr151 and hbb genes is also over 27-fold (one-way ANOVA, *p<0.05). Taken together the results suggest that OR promoters are in an inactive state within the genome of mESCs.

**Fig. 6.**
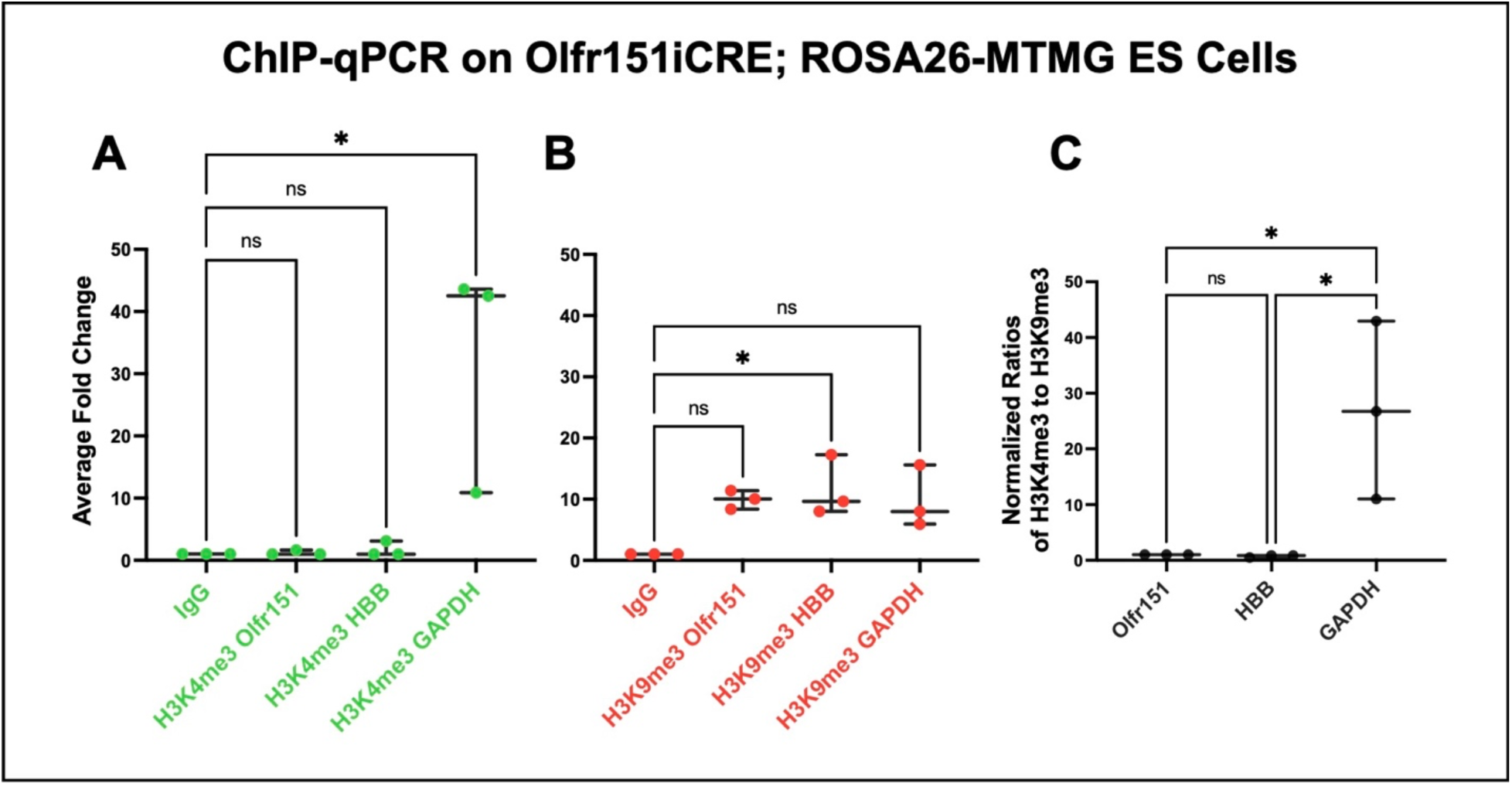
Degree of H3K4me3 and H3K9me3 epigenetic marking for Olfr151, HBB, and GAPDH. ChIP-qPCR readings for the presence of histones **(A)** H3K4me3 and **(B)** H3K9me3 on Oflr151, β-globin (HBB), and GAPDH in Olfr151iCRE; ROSA26-MTMG reporter. Raw data in Supplemental Table 3. Each dot represents a separate experiment replicate. ANOVA analysis reveals H3K9me3 HBB shows statistical significance relative to IgG control at p< 0.029, whereas Olfr151 and GAPDH significance is at p<0.065 and p<0.68 respectively. By contrast, H3K4me3 were no different than IgG control, but GAPDH is at p<0.14. **(C)** Overall, the ratios of H3K4me3 (on marks) to H3K9me3 (off marks) show GAPDH at 27-fold higher than Olfr151 and 36-fold higher than HBB, p<0.03.

### Olfr151iCRE allele in olfactory sensory neurons

All odorant receptors are expressed in a singular fashion (a single allele is expressed per neuron). The Olfr151 gene is chosen to be expressed in just a few thousand OSNs out of the 10 million total. This low frequency of expression (“choice”) could be difficult to activate in mESC model system. We have previously shown that a strong choice enhancer (x21 element) can increase the probability of choice for the Olfr151 minigene by 1000-fold in mice (*4, 10, 33*). We sought to reproduce this finding using the ROSA26-MTMG reporter mouse line (Fig. 7A). As a proof of concept, four strains were produced with 5×21-Olfr151iCRE transgenic mice (Fig. 7B-E). It has previously been shown by us and others that the size of the glomerulus correlates with the number of OSNs expressing a given OR gene in OSNs. For example, the relatively small glomeruli for Olfr151 with less than 50-micron diameters correlate to just a few thousand OSNs expressing Olfr151 (Fig. 1A,B) (*33, 34*). By contrast, previously published 5×21-Olfr151-IRES-TauCherry mice (*33*) as well as all four strains (Fig. 7B-E) were at least 200-micron in diameter, which is equivalent to a 4-fold radial increase in volume or at least 268-fold more OSNs (*34*). Thus, 5×21 enhancer provides greater accessibility to the Olfr151 promoter for expression.

**Fig. 7:**
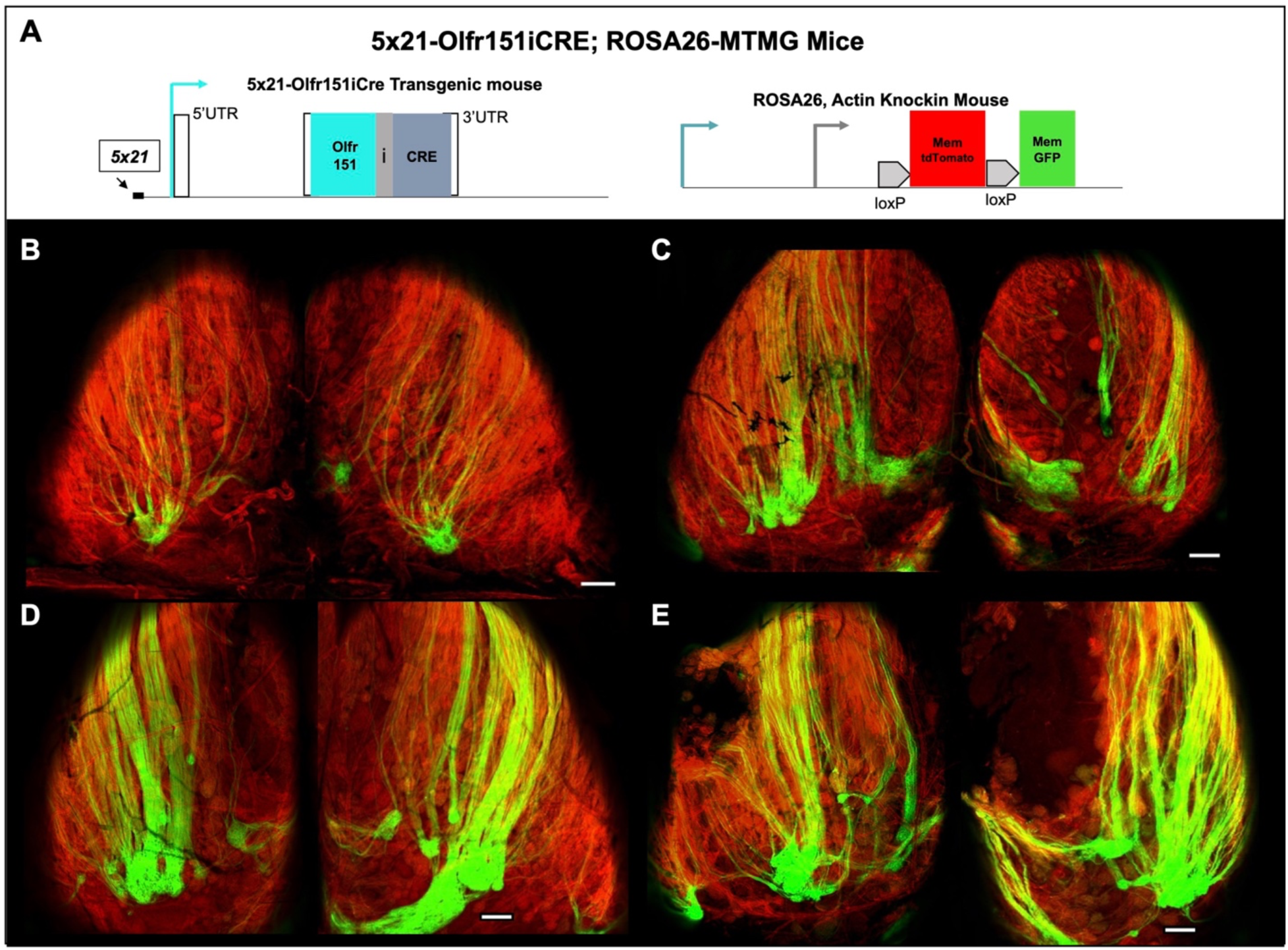
Four 5×21-Olfr151-IRES-CRE transgenes with ROSA26-MTMG reporter mouse. **(A)** Olfr151iCRE transgene mouse containing 5×21 enhancer were generated and crossed with ROSA26-MTMG mice. **(B)** 5×21-Olfr151iCre strain β (beta). **(C)** 5×21-Olfr151iCre strain ɣ (delta). **(D)** 5×21-Olfr151iCre strain ɣ (gamma). **(E)** 5×21-Olfr151iCre strain α (alpha). **(B-E)** All four transgenic lines contained coalesced GFP labeled axons into glomeruli that were larger than those found in the Olfr151-IRES-CRE knockin mice (Fig. 1B), which is due to the coalescence of greater number of Olfr151 axons. Scale bar = 200µm.

### Epigenetic Library Screens

Thus far, the Olfr151iCRE locus gene appeared resistant to expression with the tested chemical libraries. Perhaps this resistance to expression was due to the H3K9me3 marks on the gene and/or the inherent low probability of choice of the Olfr151 gene. To test the effect of the x21 enhancer on Olfr151 minigene expression in the mESC reporter line, 5 versions of the Olfr151iCRE minigene were transfected into Olfr151iCre; ROSA-MTMG ESCs (0×21, 4×21, 5×21, 7×21 and 9×21) (Fig. 8A-E) and grown for a week to allow for random integration into the genome. Only the 7×21 and 9×21 enhanced minigenes showed any CRE expression as viewed by the membrane-GFP fluorescence. Resistance to expression from 0×21, 4×21, and 5×21-Olfr151 promoter was confirmed by replacing it with the E1Fα promoter (1Kb)(*35*) and transfection into our mESC reporter line. Indeed, membrane-GFP expression was readily apparent (Fig. 8F). It should be noted that co-electroporations with a puromycin selectable marker and 5×21-Olfr151iCRE minigene still yielded no colonies with membrane-GFP expression after puromycin selection.

**Fig. 8.**
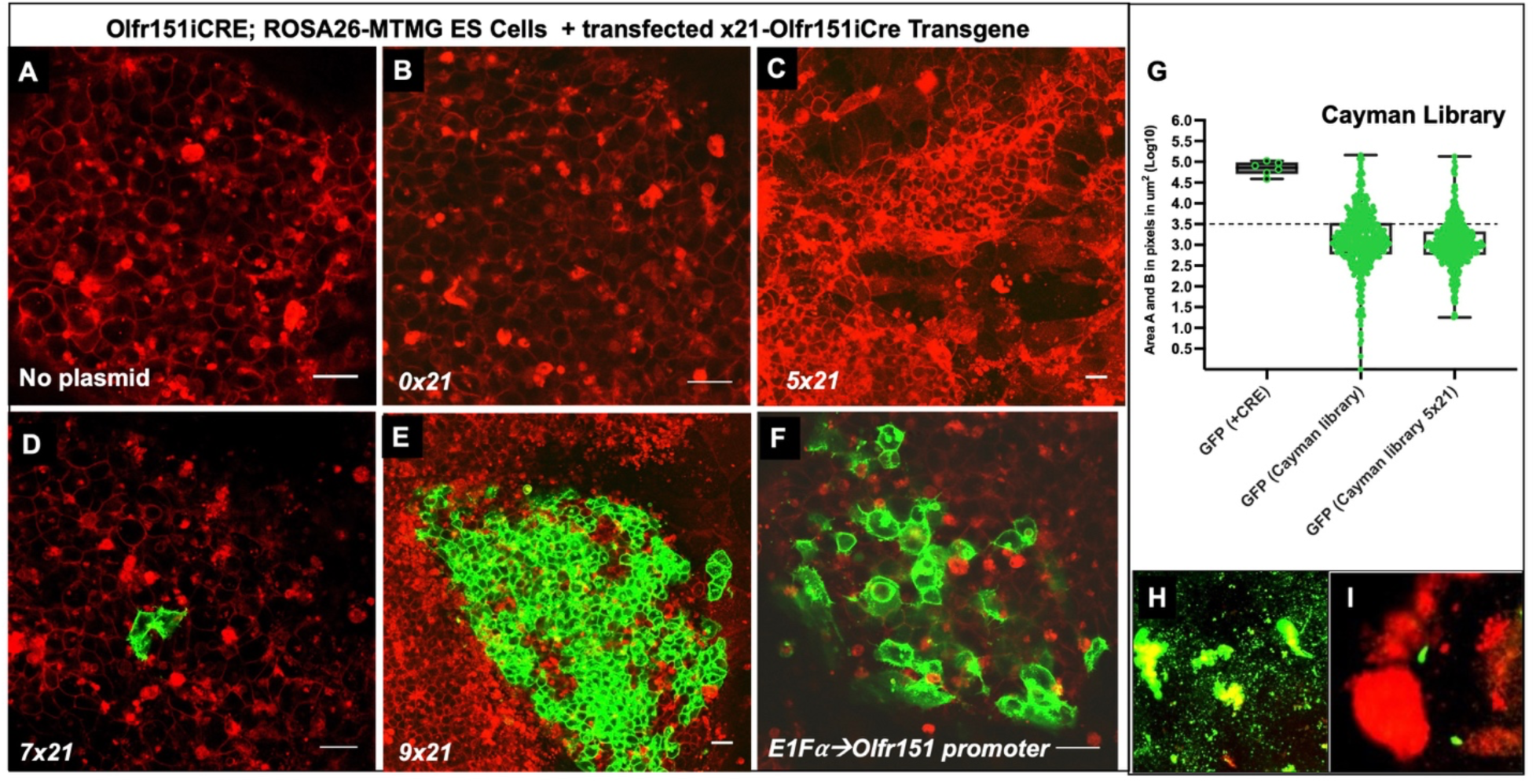
Cayman epigenetic compound treatment images of 5×21-Olfr151iCRE minigene transfections into *Olfr151iCRE; ROSA26-MTMG* reporter mESCs. **(A)** Native/untreated Olfr151iCRE; ROSA26-MTMG mESCs. (B-F) CRE-containing minigenes were transfected in Olfr151iCRE; ROSA26-MTMG reporter mESCs. **(B)** Fluorescent mages of mESCs transfected with 0×21*-*Olfr151iCRE minigene. **(C)** Fluorescent mages of mESCs transfected with 5×21*-*Olfr150iCRE minigene**. (D)** Fluorescent mages of mESCs transfected with 7×21*-*Olfr151iCRE minigene. **(E)** Fluorescent mages of mESCs transfected with 9×21*-*Olfr151-IRES-CRE minigene. **(F)** The Olfr151 promoter was substituted with the ESC-expressed E1Fα promoter in the minigene. The images were taken 24-48 hours after transfection. No membrane GFP fluorescence was observed for 0×21- and 5×21-Olfr151iCRE minigenes. The 7×21*-* Olfr151iCRE minigene showed sporadic membrane GFP expression that was readily discernible and the 9×21*-*Olfr151iCRE minigene revealed many cells with membrane GFP expression, but the CRE control revealed fluorescence in nearly all mESC colonies. Scale Bar = 20µm. The E1Fα promoter controlled for the functionality of the Olfr151 minigene. **(G)** Olfr151iCRE; ROSA26-MTMG mESCs, transfected with 5×21*-*Olfr151iCRE minigene, were treated with 175 epigenetic chemical compounds (Cayman library: small compound epigenetic library). The cells were fixed 48hours after treatment. GFP (+ CRE) log10 Area B = 4.84 ± 0.16. In native mESCs, treated with Cayman Library compounds, the GFP fluorescence value is log10 Area B Avg = 3.10 (with 101 images above the 3.5 threshold value). In the reporter mESCs transfected with 5×21-Olfr151iCRE minigene and treated with Cayman Library compounds, the GFP fluorescence value is Avg of 3.04 with 60 images above the 3.5 threshold. A close inspection of all images from mESCs treated with Cayman Library epigenetic compounds did not reveal any images with membrane GFP expression. **(H, I)** Two examples of fluorescent expression that appeared to be promising epigenetic compound candidates but did not have membrane GFP expression (see panels D-F). **(H)** Splitomicin treatment of mESCs (the compound plays a role in silencing gene expression). **(I)** Anacardic acid treatment of mESCs (an anti-inflammatory and anti-tumor compound). (H, I) Scale bar = 100µm.

Resistance to expression of the 5×21-Olfr151iCRE minigene and the knockin Olfr151iCRE locus suggested that either transcription factors were not present in mESCs and/or epigenetic marks limited gene activation. A final attempt to activate our mESC reporter line with or without a transfected 5×21-Olfr151iCRE was attempted using a specialized epigenetic library by Cayman Chemicals. This library solely includes 175 epigenetic compounds, which specifically interfere with the activity of enzymes necessary for epigenetic modifications, such as methyltransferases, demethylases, histone acetyltransferases, and histone deacetylases, which methylate, demethylate, add acetylation groups, or remove them from the marked genes (*36*). For the screening, all compounds were utilized at two concentrations, 1µM and 10µM. However, based on the lack of membrane-GFP expression, not one of these chemicals was a candidate for retesting (Fig. 8G-I).

## Discussion

The ability to rigorously study the mechanism of odorant receptor gene choice has been significantly hampered by the lack of any olfactory cell lines that foster OR expression or differentiation into post-mitotic cells that confer OR expression. Hence, this mechanism has only been studied in the olfactory epithelium of mice. Here, a mouse embryonic stem cell (mESC) reporter system was utilized to establish a one-step mechanism to trick mESCs to express an OR gene even in a few cells. Nearly 5000 chemicals tested at two different concentrations failed to elicit any leaky expression of the Olfr151iCRE allele in sufficient quantities to trigger recombination in the ROSA26-MTMG reporter (See Table 1). The presence of the epigenetic mark H3K9me3 and the absence of H3K4me9 at the Olfr151 promoter in mESCs could be contributing to why the Olfr151 promoter failed to be activated. A chemical library containing compounds that can alter the epigenetic landscape at gene promoters also failed to elicit any convincing membrane-GFP expression.

**Table 1.**
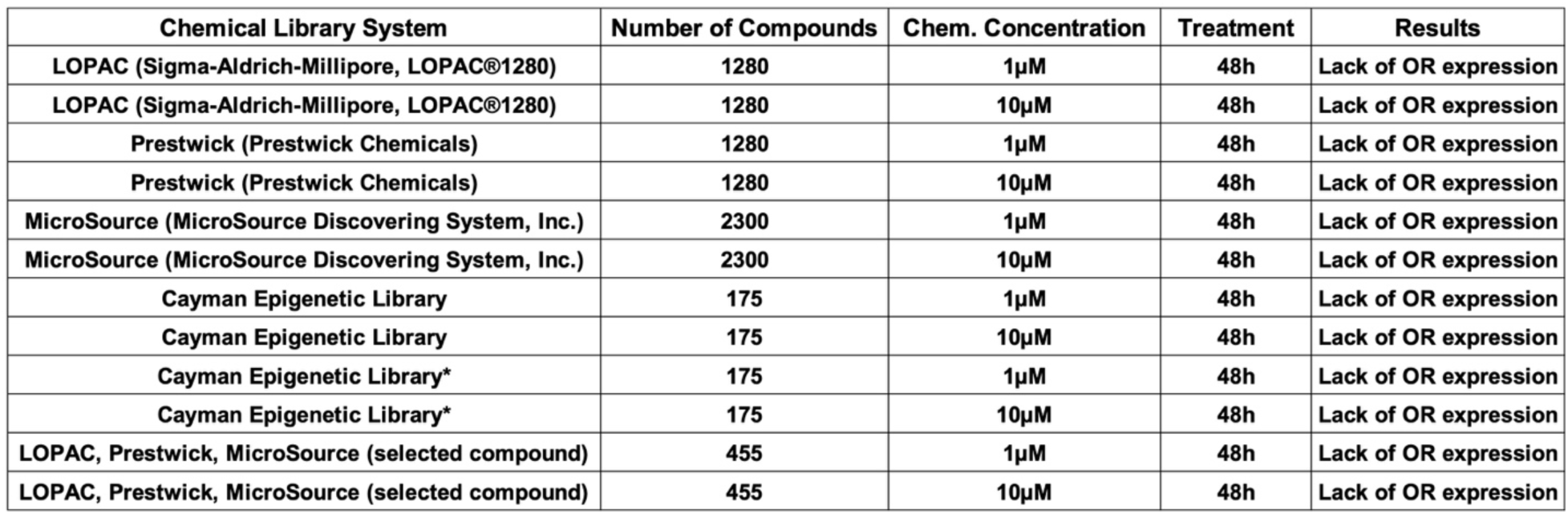
Chemical compound libraries utilized and outcomes. Chemical concentration, treatment period in the reporter system, and GFP expression. None of the compounds revealed convincing membrane GFP expression. *Screened post 5×21-Olfr151iCRE minigene transfection.

| Chemical Library System | Number of Compounds | Chem. Concentration | Treatment | Results |
| --- | --- | --- | --- | --- |
| LOPAC (Sigma-Aldrich-Millipore, LOPAC®1280) | 1280 | 1µM | 48h | Lack of OR expression |
| LOPAC (Sigma-Aldrich-Millipore, LOPAC®1280) | 1280 | 10µM | 48h | Lack of OR expression |
| Prestwick (Prestwick Chemicals) | 1280 | 1µM | 48h | Lack of OR expression |
| Prestwick (Prestwick Chemicals) | 1280 | 10µM | 48h | Lack of OR expression |
| MicroSource (MicroSource Discovering System, Inc.) | 2300 | 1µM | 48h | Lack of OR expression |
| MicroSource (MicroSource Discovering System, Inc.) | 2300 | 10µM | 48h | Lack of OR expression |
| Cayman Epigenetic Library | 175 | 1µM | 48h | Lack of OR expression |
| Cayman Epigenetic Library | 175 | 10µM | 48h | Lack of OR expression |
| Cayman Epigenetic Library* | 175 | 1µM | 48h | Lack of OR expression |
| Cayman Epigenetic Library* | 175 | 10µM | 48h | Lack of OR expression |
| LOPAC, Prestwick, MicroSource (selected compound) | 455 | 1µM | 48h | Lack of OR expression |
| LOPAC, Prestwick, MicroSource (selected compound) | 455 | 10µM | 48h | Lack of OR expression |

**Table 1**
| **Chemical Library System** | **Number of Compounds** | **Chem. Concentration** | **Treatment** | **Results** |
| --- | --- | --- | --- | --- |
| **LOPAC (Sigma-Aldrich-Millipore, LOPAC®1280)** | **1280** | **1μM** | **48h** | **Lack of OR expression** |
| **LOPAC (Sigma-Aldrich-Millipore, LOPAC®1280)** | **1280** | **10μM** | **48h** | **Lack of OR expression** |
| **Prestwick (Prestwick Chemicals)** | **1280** | **1μM** | **48h** | **Lack of OR expression** |
| **Prestwick (Prestwick Chemicals)** | **1280** | **10μM** | **48h** | **Lack of OR expression** |
| **MicroSource (MicroSource Discovering System, Inc.)** | **2300** | **1μM** | **48h** | **Lack of OR expression** |
| **MicroSource (MicroSource Discovering System, Inc.)** | **2300** | **10μM** | **48h** | **Lack of OR expression** |
| **Cayman Epigenetic Library** | **175** | **1μM** | **48h** | **Lack of OR expression** |
| **Cayman Epigenetic Library** | **175** | **10μM** | **48h** | **Lack of OR expression** |
| **Cayman Epigenetic Library\*** | **175** | **1μM** | **48h** | **Lack of OR expression** |
| **Cayman Epigenetic Library\*** | **175** | **10μM** | **48h** | **Lack of OR expression** |
| **LOPAC, Prestwick, MicroSource (selected compound)** | **455** | **1μM** | **48h** | **Lack of OR expression** |
| **LOPAC, Prestwick, MicroSource (selected compound)** | **455** | **10μM** | **48h** | **Lack of OR expression** |

Furthermore, transfection of the 5×21-Olfr151iCRE minigene in the mESCs did not show position effect variegation to produce membrane-GFP expression and would only be subject to proximal epigenetic states of the chromatin. This was also true when 5×21-Olfr151iCRE transfection was subjected to antibiotic selection. Finally, applying the Cayman chemical library to x21-minigene-transfected mESCs also failed to elicit membrane-GFP expression. Conversely, the substitution of the Olfr151 promoter with an E1Fα promoter, known to be expressed in ESCs, yielded colonies with membrane-GFP expression (Fig. 8F), suggesting that the Olfr151 promoter is kept silent and/or there is a lack of transcription factors in mESCs that can lead to Olfr151 promoter activity.

The Olfr151 promoter contains two recognizable conserved DNA sequences that likely bind the transcription factors LIM homeodomain protein, LHX2 and the Olf1/Ebf1 (O/E) family of proteins (O/E1-4), neither of which is expressed in mESCs (*22, 37–39*). That said, O/E1 is expressed by B-cells, but no OR gene expression has been reported in these cells. By contrast, LHX2 is utilized for neural development, yet OR expression remains the province of the olfactory sensory neurons. Interestingly, all four of the 5×21-Olfr151iCRE transgenes generated very large (at least 4x the radius wildtype) membrane-GFP labeled glomeruli along with their axonal tracks, with a paucity of any other labeled axonal tracks or innervation of other glomeruli throughout the olfactory bulb (Fig. S3B-E). Only the 5×21-Olfr151iCREα strain revealed some GFP expression in random glomeruli (Fig. S3E). In comparison, only sporadic leaky OSN expression is observed in the 7×21 enhancer-driven Olfr151iCRE minigenes within mESCs, suggesting tight regulation of the OR choice mechanism. These results lend support to the notion that 5×21 minigenes are not leaky in early OSN progenitors or postmitotic OSNs at sufficient levels to produce enough CRE protein to recombine the ROSA26-MTMG reporter.

The early occurrence of negative epigenetic marks on ORs might result from their location in OR clusters (approximately 30 OR clusters exist in the mouse genome). The presence of ORs in gene arrays could be causative for this negative repression. It has been shown that the β2 adrenergic receptor has no nucleotide homology to OR coding sequences, substitutes for Olfr151 receptor in knock-in experiments, and follows the same mechanism of OR gene choice. These results suggest that the coding region DNA does not contribute to negative repression (*8, 9, 40*). In addition, no homology exists in the 5’UTRs, introns, or 3’UTRs across all OR genes. Thus, the silencing of OR promoters may be a feature of the conserved and paired LHX2 and O/E binding sites. These paired sites are found in the Greek Island super enhancers in all OR clusters (*17*).

A future experiment could include a high-throughput chemical screen on an OSN precursor cell line containing OR dependent CRE expression and CRE-reporter genes. The olfactory placodal cell line, OP6 (*29*), could be such a candidate. In addition, a cocktail of small chemical compounds could be tested; unfortunately, this greatly increases the possible combinations. If the OR expression could be reliably produced in vitro in the future, then it could prove to be a reliable platform to characterize the mechanism of OR expression.

### Conclusion

The main goal of the discussed experiments was to create a one-step method to characterize the promoter activation of odorant receptors. The inability of Olfr151 activation in mESCs and the lack of leaky expression in mitotic OSN precursors suggests a highly organized mechanism of OR expression specific to postmitotic olfactory sensory neurons.

## Acknowledgements

Special thanks to Shuibing Chen at Weill Cornell Medical Institute for providing logistical and intellectual support that helped drive the success of these experiments. This work was supported by the NYSTEM C023048 grant, NIH SC1 GM088114 and NIH R01 DC020764. We thank the Hunter College Animal Facility Manager, Barbara Wolin, and Veterinarian Patricia Glennon for help in maintaining the transgenic colony, and Rada Novinsky at the Transgenic Core Facility at The Rockefeller University for generating transgenic founders.

All data generated are found in the supplementary files of this manuscript.

---

**Supplementary Fig. S1.**
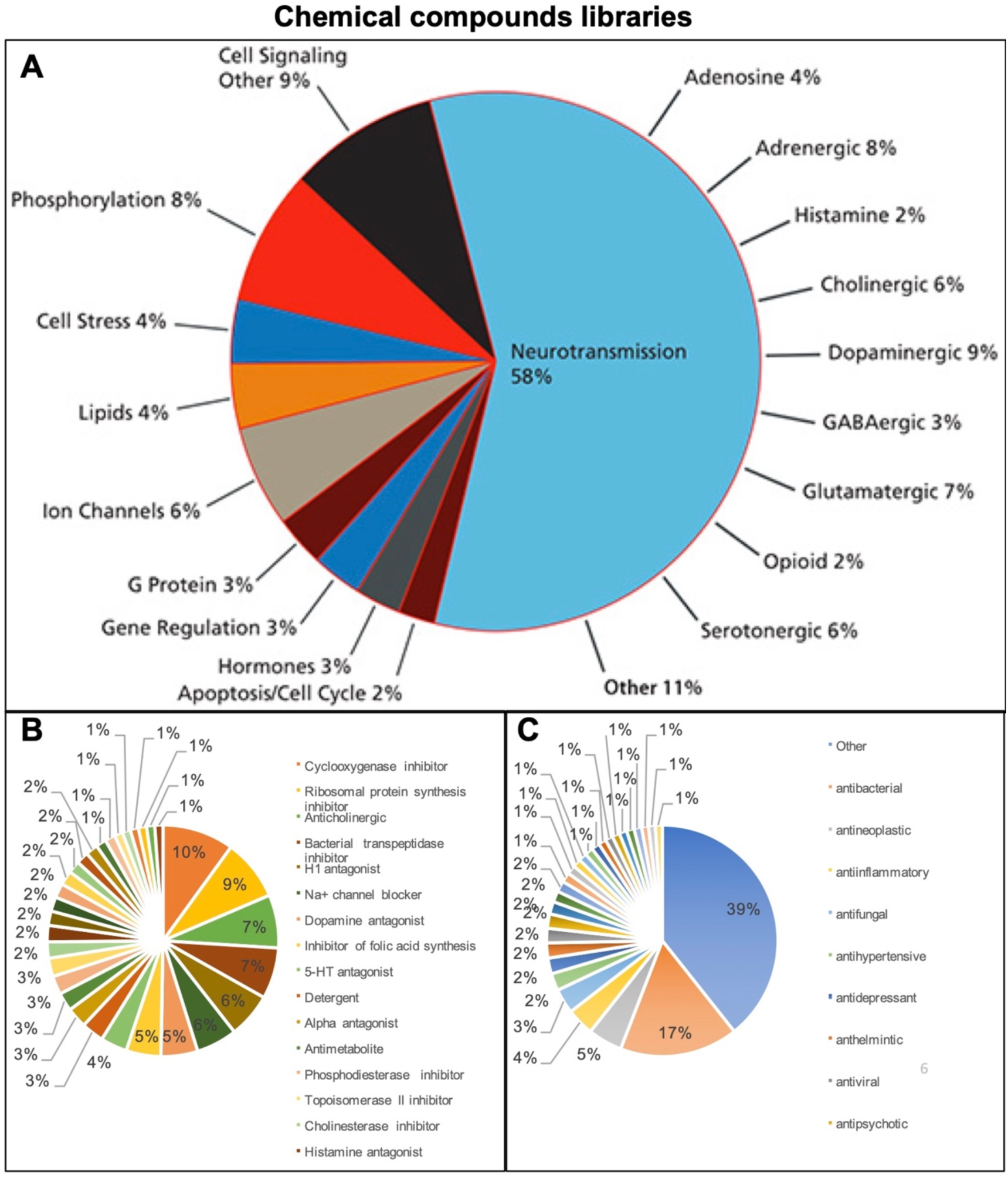
Chemical compounds libraries. **(A)** 1280 LOPAC chemical compounds are grouped together based on the mechanisms of their activities (LOPAC^®1280^). **(B)** Prestwick chemical compounds are grouped based on their chemical activity (Prestwick Chemical). **(C)** MicroSource chemical compounds are grouped based on similar mechanisms of activity upon the cells (MicroSource Discovery Systems, Inc.).

**Supplementary Fig. S2.**
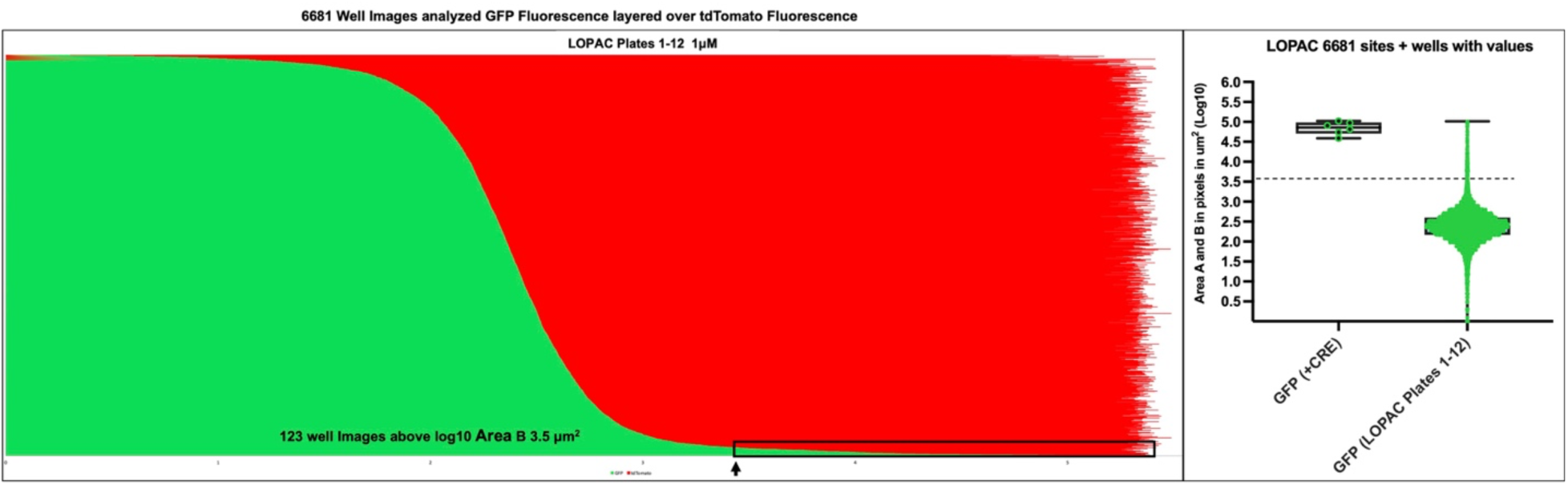
Fluorescent expression in mESCs as a result of LOPAC chemical compounds treatments. All LOPAC treatments (12x 384-well plates, in two concentrations) yielded 6881 images (A combination of nine and four images per well) log10 Area (fluorescence area, µm^2^) of tdTomato (red) and GFP (green) fluorescence in reporter cells. The green and red log10 Area values were ranked based on increasing green fluorescence (log10 Area B, µm^2^). 123 images contained GFP fluorescence shown within black rectangle above the threshold of log10 Area B = 3.5µm^2^ (see Fig. 4).

**Supplementary Fig. S3.**
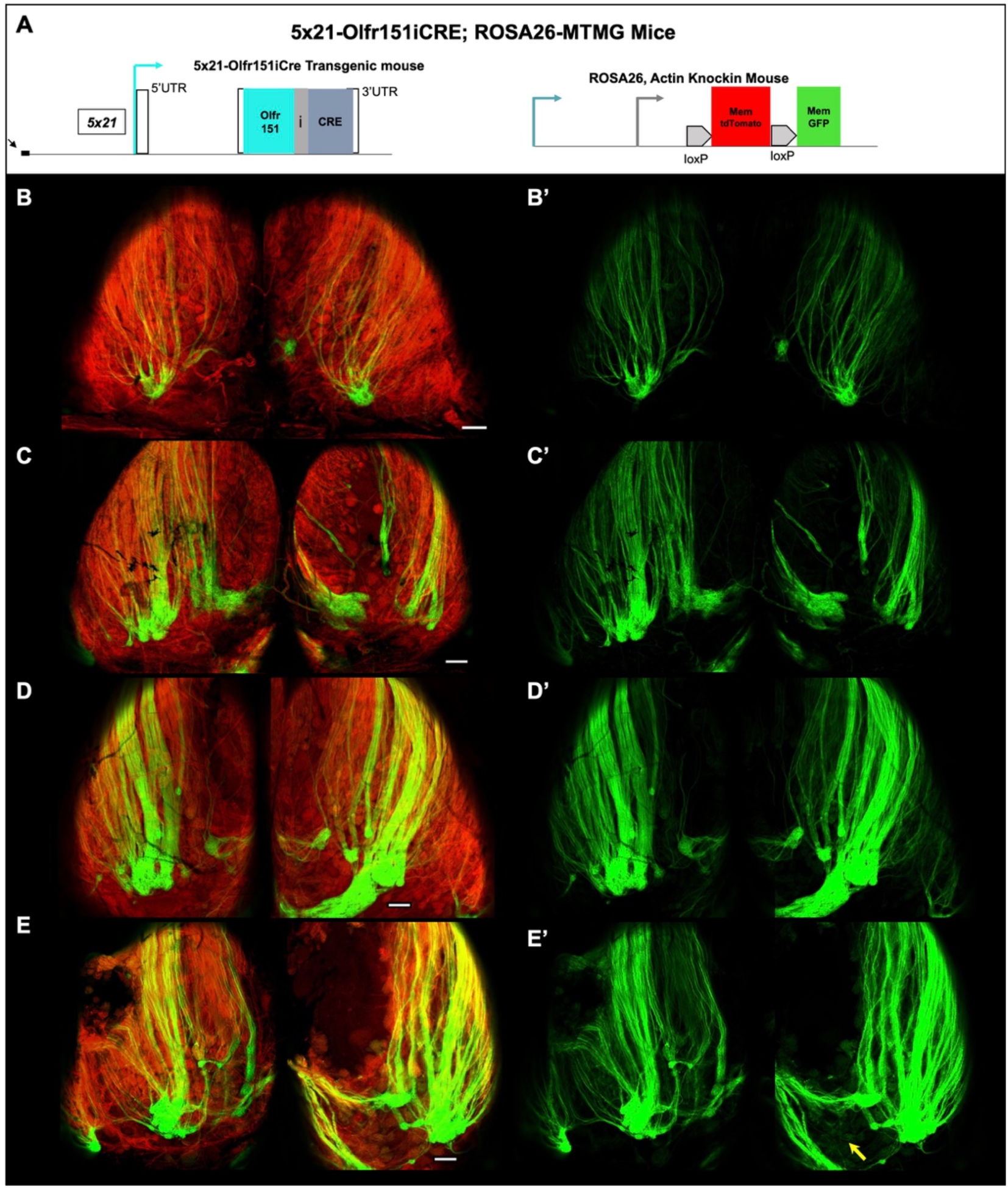
Four 5×21-Olfr151iCRE transgenes in the ROSA26-MTMG reporter mouse. **(A-D)** Overlays of wholemount tdTomato and GFP fluorescent microscopy of olfactory bulbs. (A) Olfr151iCRE transgene mice, containing 5×21 enhancer, were generated and crossed with ROSA26-MTMG mice. (B) 5×21-Olfr151iCre strain β (beta) line. (C) 5×21-Olfr151iCre strain ɣ (delta). (D) 5×21-Olfr151iCre strain ɣ (gamma). (E) 5×21-Olfr151iCre strain α (alpha). (B-E) All four transgenic lines contained coalesced GFP labeled axons into glomeruli that were larger than those found in the Olfr151iCRE knockin mice (Fig. 1B), which is due to the coalescence of greater number of Olfr151 axons. Scale bar = 200µm. **(A’-D’)** Wholemount GFP fluorescent microscopy of olfactory bulbs from (A-D).

